# Advancing thermostability of the key photorespiratory enzyme glycerate 3-kinase by structure-based recombination

**DOI:** 10.1101/2024.05.02.592181

**Authors:** Ludmila V. Roze, Anna Antoniak, Daipayan Sarkar, Aaron H. Liepman, Mauricio Tejera-Nieves, Josh V. Vermaas, Berkley J. Walker

## Abstract

As global temperatures rise, maintaining and improving crop yields will require enhancing the thermotolerance of crops. One approach for improving thermotolerance is using bioengineering to increase the thermostability of enzymes catalyzing essential biological processes. Photorespiration is an essential recycling process in plants that is integral to photosynthesis and crop growth. The enzymes of photorespiration are targets for enhancing plant thermotolerance as this pathway limits carbon fixation at elevated temperatures. Exploring inter-specific variation of the key photorespiratory enzyme glycerate kinase (GLYK) from various photosynthetic organisms, we found that the homolog from the thermophilic alga *Cyanidioschyzon merolae* was more thermotolerant than those from mesophilic plants, including *Arabidopsis thaliana*. To understand factors influencing thermotolerance of *C. merolae* GLYK (CmGLYK), we performed molecular dynamics simulations using AlphaFold-predicted structures, which revealed greater movement of loop regions of mesophilic plant GLYKs at higher temperatures compared to CmGLYK. Based on these simulations, a series of hybrid proteins were produced and analyzed. These hybrid enzymes contained selected loop regions from CmGLYK replacing the most highly mobile corresponding loops of AtGLYK. Two of these hybrid enzymes had enhanced thermostability, with melting temperatures increased by 6 °C. One hybrid with three grafted loops maintained higher activity at elevated temperatures. While this hybrid enzyme exhibited enhanced thermostability and a similar K_m_ for ATP compared to AtGLYK, its K_m_ for glycerate increased threefold. This study demonstrates that molecular dynamics simulation-guided structure-based recombination offers a promising strategy for enhancing thermostability of other plant enzymes.

## Introduction

Global temperatures are rising at a rapid rate and will continue to rise for the foreseeable future, vastly impacting many processes important for humans like plant productivity (Calvin *et al*., 2023; Ditlevsen and Ditlevsen, 2023; Doughty *et al*., 2023). Elevated temperature affects many aspects of plant growth and development, which can decrease crop yield. Much effort has been directed at improving the temperature response of plants by targeting specific genes, hormone signaling pathways, and epigenetic regulation (Ashraf, 2021; Chen *et al*., 2022; Moore *et al*., 2021) and the resulting temperature-dependent decline in the function of the photosynthetic apparatus and cellular metabolism which power plant growth and shape yield (Wahid *et al*., 2007). Therefore, engineering improved thermotolerance in photosynthesis and related metabolic pathways into crop plants for improved productivity in warmer temperatures may help sustain yields as global temperatures rise.

Rubisco is the enzyme that catalyzes CO_2_ fixation and is also responsible for much of the temperature sensitivity of photosynthesis. This temperature sensitivity comes in part since rubisco also binds oxygen, and when it does, the subsequent oxygenation reaction produces the inhibitory metabolite 2-phosphoglycolate. Photorespiration is the metabolic pathway that recycles 2-phosphoglycolate, yielding 3-phosphoglycerate (3-PGA), which can enter the C_3_ cycle (Bauwe, 2023; Somerville, 2001). While photorespiration is necessary, the pathway consumes 30-40% of C3 leaf energy and 25-30% of carbon fixed by rubisco is released via this pathway as a result of its glycine decarboxylase activity (Bauwe, 2023; Walker *et al*., 2016). At elevated temperatures, rubisco’s affinity for CO_2_ and the solubility of CO_2_ (relative to O_2_) decrease, increasing relative rates of rubisco oxygenation and photorespiration (Bernacchi *et al*., 2002; Jordan and Ogren, 1984; Walker *et al*., 2016, 2013). Aside from its integral connection with photosynthesis, photorespiration also interfaces with key pathways of primary metabolism, including nitrogen assimilation, one-carbon metabolism, and protein synthesis (Bauwe *et al*., 2012; Fu and Walker, 2023; Hanson and Roje, 2001). Given its importance supporting photosynthesis under elevated temperatures and contributing to other metabolic pathways, photorespiration therefore poses an attractive bioengineering target.

Examples from nature and engineering experiments indicate that photorespiratory capacity may become limited at elevated temperatures and that it may be possible to optimize enzyme activity to maintain photorespiration at elevated temperatures. For example, in *Rhazya stricta*, a plant that grows at high temperatures (43-52 °C) in the deserts of Saudi Arabia and Pakistan, high rates of CO_2_ assimilation at elevated temperature are associated with high rates of photorespiration that correlate with increased activity of photorespiratory enzymes (Gregory *et al*., 2023). Additionally, increasing the activity of certain photorespiratory enzymes or introducing a synthetic photorespiratory pathway that metabolizes glycolate in chloroplasts increases net CO_2_ fixation at elevated temperatures, supporting the hypothesis that photorespiratory capacity can be limiting under elevated temperature and improved through engineering (Timm *et al*., 2019; Cavanagh *et al*., 2022; Kebeish *et al*., 2007; South *et al*., 2019). These studies provide evidence that it may be possible to engineer thermotolerance in crops with increased photosynthetic activity by maintaining the stability and activity of photorespiratory enzymes at elevated temperatures.

D-glycerate 3-kinase (EC 2.7.1.31; GLYK) is a chloroplastic enzyme that catalyzes the final step of photorespiration, converting glycerate into 3-PGA (Figure 1) that can enter the C_3_ cycle (Boldt *et al*., 2005). A single *GLYK* gene (AT1G80380) is present in the *Arabidopsis thaliana* genome. The activity of *A. thaliana* GLYK (AtGLYK) is essential for viability of plants grown in ambient air, though knockout mutant plants can grow under nonphotorespiratory conditions of elevated CO_2_ (Boldt *et al*., 2005). Because GLYK activity is imperative for plant growth and development under ambient atmospheres, inactivation or insufficient thermotolerance of this enzyme due to heat could limit plant productivity, making this enzyme an attractive target for engineering increased thermotolerance.

**Figure 1.**
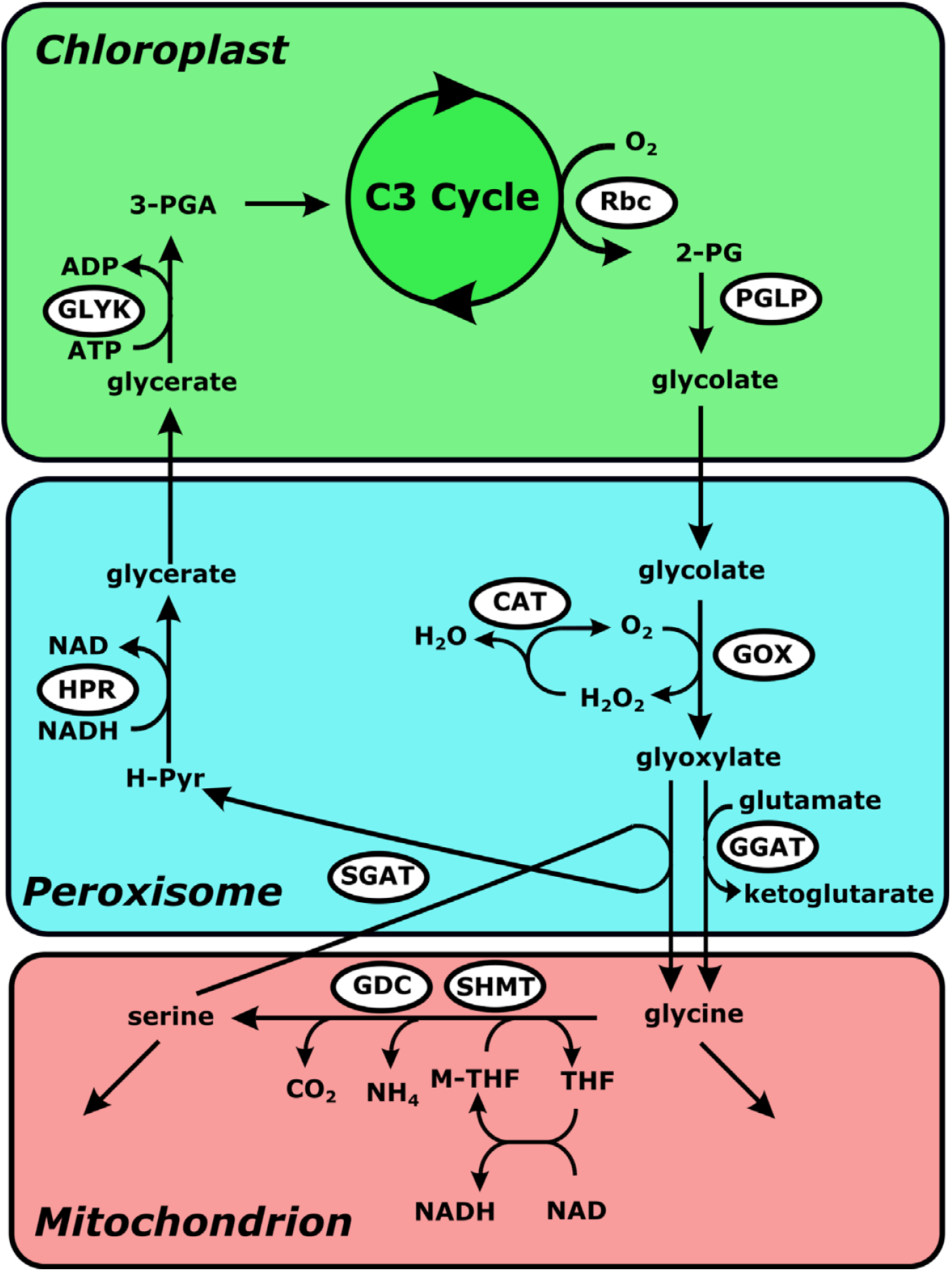
The photorespiratory pathway. Photorespiratory enzymes are located in closely spaced cellular organelles comprising chloroplasts, peroxisomes, and mitochondria. The pathway starts in the chloroplasts with the binding of rubisco of oxygen to Ribulose 1,5 bisphosphate to form 2-phosphoglycolate (2-PG). 2-PG is recycled to 3-phosphoglycerate (3-PGA) which enters the C3 cycle. Abbreviations include: Rbc, rubisco; 2-PG, 2-phosphoglycolate; PGLP, phosphoglycolate phosphatase; GOX, glycolate oxidase; CAT, catalase; GGAT, glutamate:glyoxylate aminotransferase; SGAT, serine:glyoxylate aminotransferase; HPR, hydroxypyruvate reductase; SHMT, serine hydroxymethyltransferase; GDC, glycine decarboxylase complex; THF, tetrahydrofolate; M-THF, methyl- tetrahydrofolate; H-Pyr, hydroxypyruvate; GLYK, glycerate kinase; 3-PGA, 3- phosphoglyceric acid.

Several approaches have been used for increasing enzyme thermotolerance generally and plant enzymes specifically. Previously, the thermostability of rubisco activase was enhanced by random recombination of gene fragments and targeted mutations (Kurek *et al*., 2007; Scafaro *et al*., 2019). While successful in this case, random recombination can generate large numbers of possible thermotolerant enzymes to analyze, potentially limiting the use of this approach.

Another approach for increasing protein thermostability while maintaining enzymatic activity involves manipulating structural enzyme loops, also termed “structure-based recombination” (Tawfik, 2006; Yedavalli and Rao, 2013; Yu *et al*., 2017; Zheng *et al*., 2019). Loops are vital elements of protein structures and affect many aspects of protein function, including substrate recognition and specificity, signal transduction, protein-protein interaction, and thermostability (Corbella *et al*., 2023; Feller and Lewitzky, 2012; Yoshida *et al*., 2021; Zheng *et al*., 2019).

However, loop regions and their conformational dynamics at elevated temperatures are difficult to predict from the sequence alone, hence an ensemble representation is invaluable for determining them *in silico* (Nussinov *et al*., 2023; Sarkar *et al*., 2023). Using protein structure prediction tools such as AlphaFold, a protein sequence can be expanded into a three-dimensional protein structure and loop regions identified. The static snapshot of the enzyme that AlphaFold provides can then be dynamically explored by classical molecular dynamic simulation to simulate protein conformational change at elevated temperature.

Here we applied a novel approach to inform structure-based recombination by using molecular dynamics simulations of AlphaFold-generated structural models. Specifically, we aimed to enhance the thermostability of AtGLYK by introducing specific corresponding segments from a more thermostable GLYK homolog from *Cyanidioschyzon merolae* (CmGLYK). *C. merolae* is a thermophilic red algal species adapted to growth in sulfuric acid hot springs (35-56 °C) (Miyagishima and Tanaka, 2021), providing an excellent template to create hybrid enzymes with improved heat tolerance. First, we compared kinetic characteristics of recombinant GLYK enzymes from *C. merolae* and several mesophilic plants (*Arabidopsis thaliana*, *Nicotiana tabacum*, *Brassica rapa*) with CmGLYK being the most thermostable. We next identified highly mobile loop regions of AtGLYK using AlphaFold-assisted molecular dynamics simulations and created hybrids by replacing them with corresponding loop sequences from the more thermostable CmGLYK. Some of the GLYK hybrids were more thermostable, demonstrating that using AlphaFold-assisted molecular dynamics simulations to inform structure-based recombination is a powerful strategy for improving thermostability of plant enzymes and a useful tool for protein engineering more generally.

## Results

### Thermostability of GLYK proteins from various species varies

The thermostability of GLYK proteins from mesophilic plants (*A. thaliana*, *Nicotiana tabacum*, *Brassica rapa*) and GLYK from a thermophilic alga (*C. merolae*) were compared using enzyme assays. One approach involved incubating purified recombinant GLYK proteins at different temperatures for varying durations, followed by continuous measurement of enzyme activity using a coupled spectrophotometric assay (Gregory *et al*., 2023). After incubation at 45 °C for 1 h, CmGLYK activity increased by 25%, while GLYK proteins from mesophilic plants lost their activity rapidly, likely due to irreversible denaturation. For example, activity fell to 50% in AtGLYK within 20 minutes at 45 °C and to undetectable values in *B. rapa* GLYK (BrGLYK-1) under those same conditions (Figure 2A).

**Figure 2.**
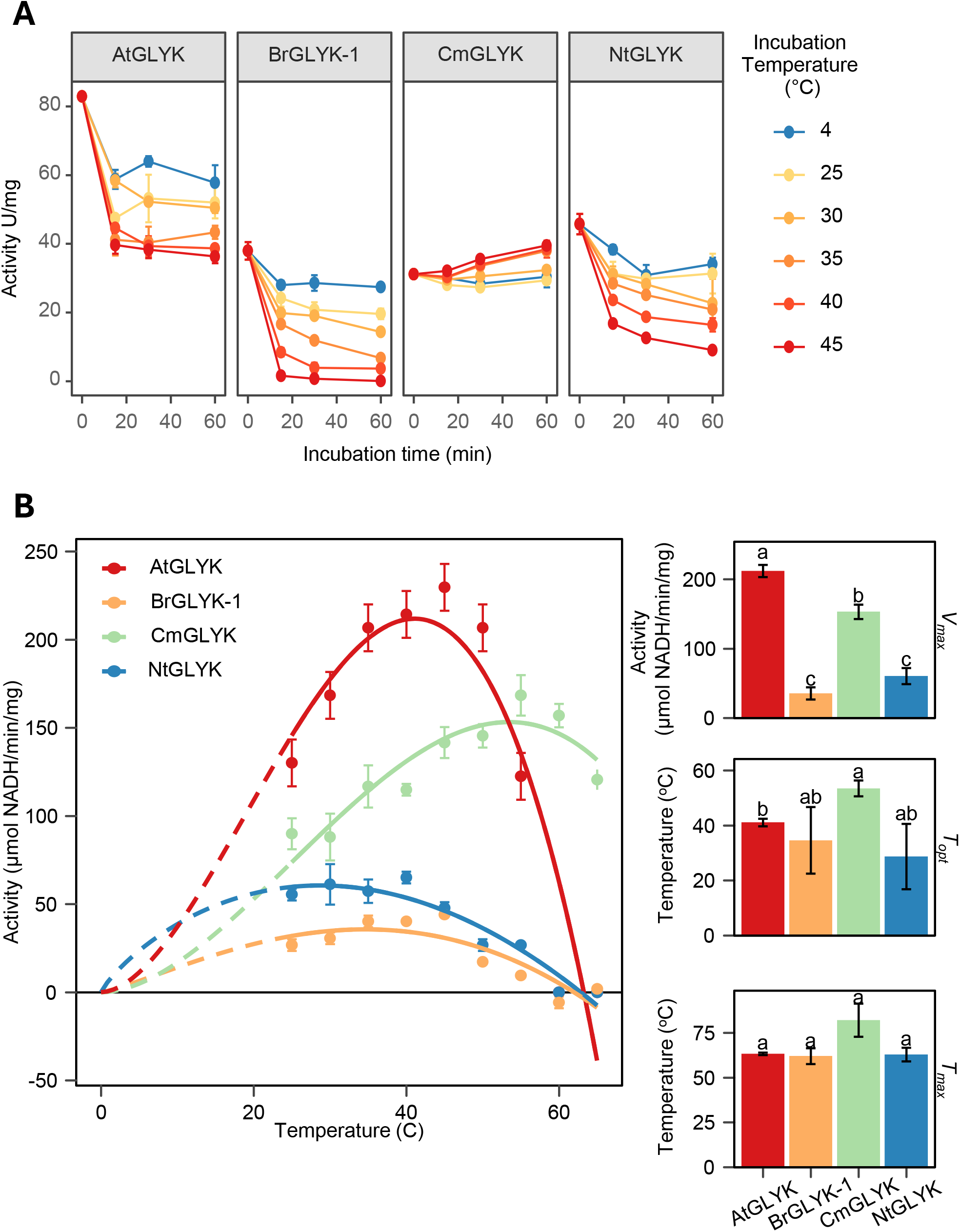
Thermal properties of glycerate kinase (GLYK) from *A. thaliana*, *B. rapa*, *C. merolae*, and *N. tabacum*. A) GLYK activity thermostability assessed by a coupled continuous spectrophotometric assay. Purified enzyme aliquots were incubated without substrates for 0, 15, 30, or 60 minutes at 25, 30, 35, 40 or 45 °C, or on ice (4°C) for control samples. Following incubation, samples were immediately assayed using the standard GLYK assay, conducted at 30 °C, as described in Experimental procedures. B) The temperature activity profile for GLYK in each species. A discontinuous 2-step assay modified from (Kehrer *et al*., 2007) was used to monitor activity of GLYK at temperatures from 25 °C to 65 °C as described in Experimental procedures. Also shown are various parameters determined from the non-linear fit of the temperature response including the maximum activity measured (V_max_), the temperature where V_max_ was measured (T_opt_), and the maximum temperature where activity would be expected (T_max_). Different lowercase letters indicate significant differences between parameters (p-value < 0.05).

In a second approach, discontinuous assays were used to measure glycerate kinase activity after incubating purified recombinant GLYK proteins with substrates at temperatures ranging between 25 °C and 65 °C. The use of discontinuous assays was necessary because the coupling enzymes used for continuous spectrophotometric assays of GLYK activity lose activity above 45°C. Discontinuous assay results indicated that optimal enzyme activity for CmGLYK occurred at 55 °C, 40 °C for AtGLYk, 30 °C for *N. tabacum* (NtGLYK), and 35 °C for BrGLYK- 1 (Figure 2B). At 60-65 °C none of the plant GLYKs were active, but CmGLYK retained approximately 80-90% of its maximal activity. Taken together, the results of both types of assays demonstrated that CmGLYK exhibited higher thermostability than plant GLYK proteins.

### Multiple sequence alignment and amino acid composition analysis revealed key differences in *C. merolae* GLYK at the level of protein primary structure

To evaluate sequence similarity among various GLYK proteins and to determine molecular characteristics that may be involved in their differential thermostability, we compared their amino acid sequences after alignment using Jalview (Troshin *et al*., 2018) (Supporting Information Figure 1). The sequence alignment showed that GLYK proteins from *A. thaliana*, *B. rapa,* and *N. tabacum* shared high sequence identity (∼90%) and similarity (∼89-97%) with one another, whereas CmGLYK had significantly lower identity (∼35-37%) and similarity (∼52-53%) to the plant enzymes. The conserved Walker A motif (G/AXXXXGKT/S) involved in ATP binding was in the N-terminal portion of all GLYK sequences (DelToro *et al*., 2016; Walker *et al*., 1982). The Walker A motif was well-conserved among plant GLYKs with the sequence APQGCGKT, while there were two amino acid deviations from the plant motif present in CmGLYK (CPQGGGKT).

A comparison of amino acid composition of the GLYK enzymes revealed a higher proportion of arginine and lower proportion of isoluecine in CmGLYK compared with GLYK enzymes from mesophilic plants (Table 1), consistent with the amino acid composition trends observed in enzymes from thermophilic bacteria, archaea and fungi (de Oliveira *et al*., 2018; Vieille and Zeikus, 2001). These trends between amino acid composition and thermostability may come from stronger intermolecular forces, such as hydrophobic packing or salt bridges, that can stabilize secondary structure at elevated temperatures. The first 90 N-terminal amino acids of CmGLYK comprise a highly variable arginine-rich region containing eight arginine residues.

**Table 1.**
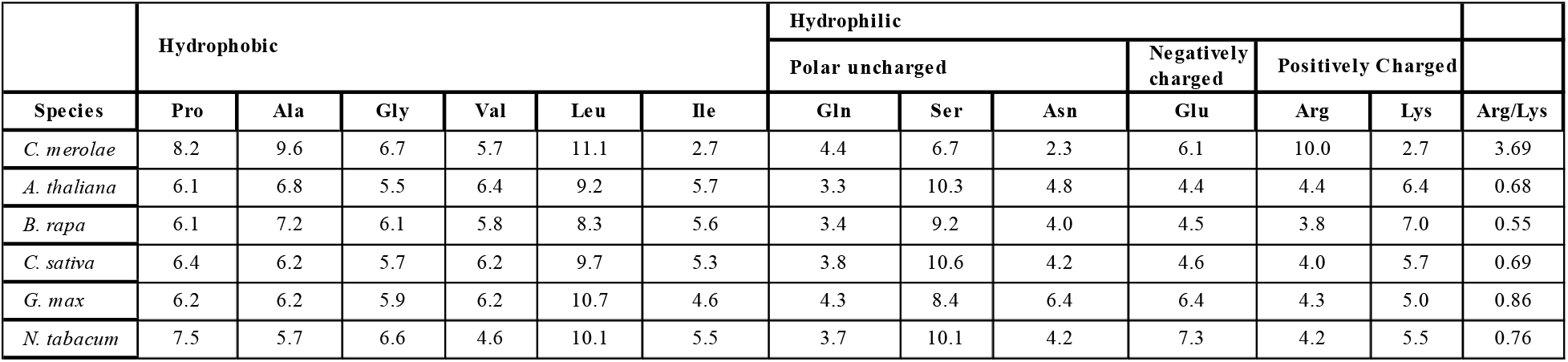
Percent of amino acid composition in various glycerate kinase (GLYK) enzymes. GLYK enzymes include a thermophilic red alga (*Cyanidioschyzon merolae*) and various plants (*Arabidopsis thaliana, Brassica rapa, Glycine max* and *Nicotiana tabacum*).

The corresponding region of AtGLYK and NtGLYK comprises three arginine residues, and in BrGLYK just two. These amino acid substitutions, and larger modifications to the overall amino acid composition may be related to the increased thermostability of CmGLYK.

### AlphaFold structure prediction and molecular simulations

To compare the predicted structures and to enable molecular dynamics simulations of GLYK proteins, we used AlphaFold (Jumper *et al*., 2021) to generate atomically detailed structural models (Supporting Information Figure 2, Animation 1). Even though CmGLYK had many variations in amino acid sequence compared to mesophilic GLYK enzymes, structurally these enzymes were similar, as expected, based on their shared catalytic activity. Using classical molecular dynamics simulations, structures for each species were solvated in a water box and simulated to create a set of trajectory snapshots at various temperatures (25, 45 and 65 **°**C, Animations 2 and 3). The enzyme dynamics within the molecular dynamics trajectories facilitate identification of protein regions that contribute to thermostability.

Determination of root-mean square fluctuations (RMSF), which quantify the movement of amino acids during the molecular dynamic simulation across temperatures (25, 45, 65 °C), for GLYK from each species revealed that three apparent surface regions of GLYK enzymes from mesophilic plants had larger RMSF at higher temperatures compared to CmGLYK (Figure 3A).

**Figure 3.**
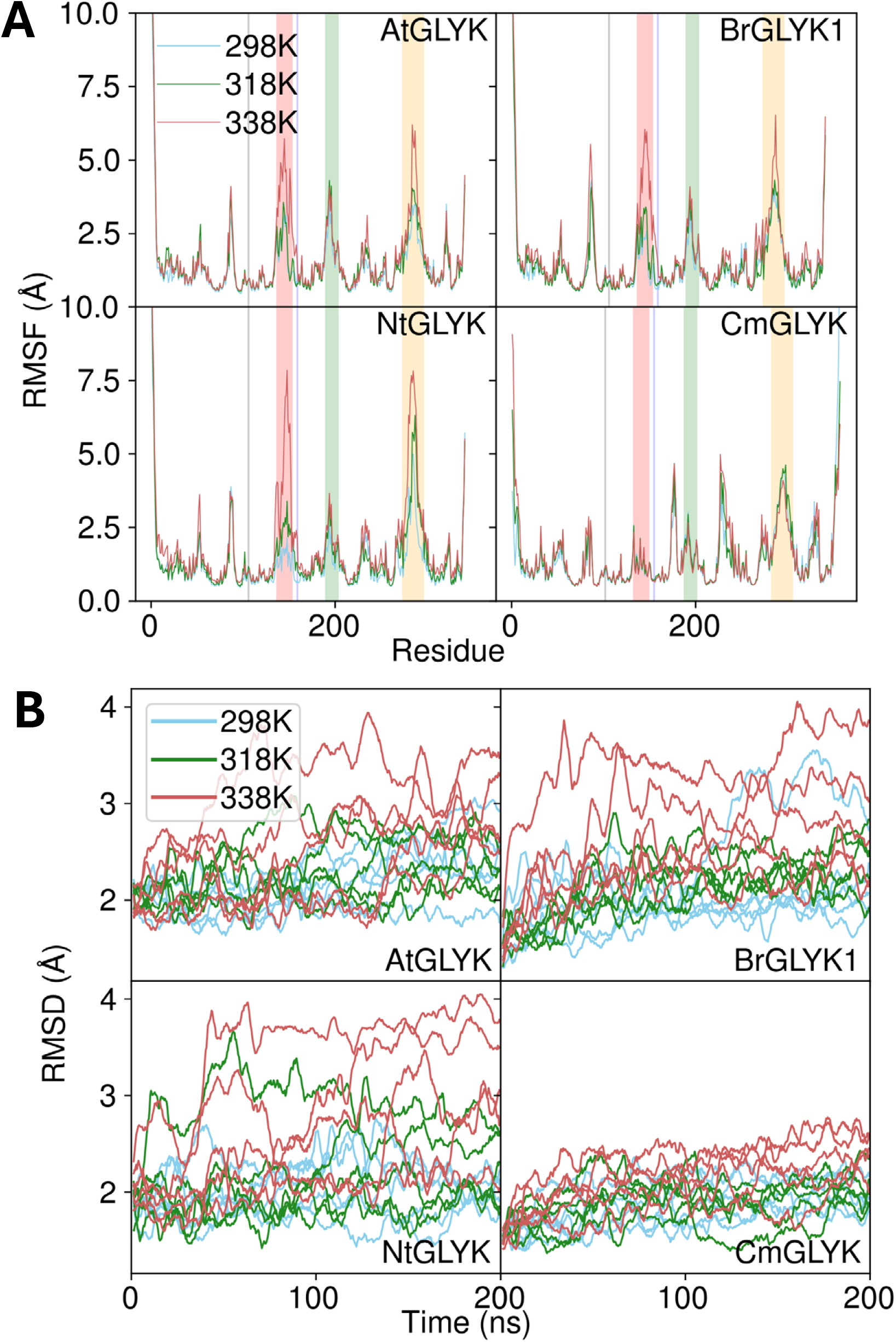
Molecular dynamics simulations for glycerate kinase (GLYK) from *A. thaliana*, *B. rapa*, *C. merolae*, and *N. tabacum*. A) Measuring fluctuations across temperatures (RMSF) in GLYK from each species with more flexible loop regions targeted in this study indicated by various shaded regions, using the same color scheme as Figure 4a. RMSF is shown relative to aligned amino acid residues across each species. B) Deviations from a reference state in time (RMSD) in the entire GLYK from each species shown over the time-dependent motions of the structure.

The fourth region, the N-terminal 90 amino acid region rich in Arg, in CmGLYK was more rigid, with a lower RMSF values in comparison to the equivalent regions from plant enzymes.

Root mean square deviation (RMSD), which quantifies the positional deviation of the protein backbone over time during the simulation, was generally between 2 to 3 Å at 25 °C (298 K) across all GLYKs. At the higher temperatures of 45 °C (318 K) and 65 °C (338 K), RMSD values for plant GLYKs tended to be higher, up to 4 Å, which indicates a protein exploring more of its configuration space, including states that may lead to unfolding over long timescales.

However, CmGLYK, which originated from a thermotolerant species *C. merolae,* had the smallest RMSD (below 3 Å) at all temperatures (Figure 3B). While the mesophile GLYK RMSDs were somewhat higher, particularly at elevated temperature, over the time course of these molecular dynamics simulations the overall protein fold remained intact in all cases.

### Design of recombinant hybrid enzymes

Molecular dynamics simulations identified three loop regions in mesophilic GLYK homologs with elevated RMSF relative to CmGLYK. One loop (Loop 3) is predicted to be in close structural proximity to the Walker A motif and the putative substrate binding site while the others are further away (Loops 1 and 2, Figure 4A). Among GLYK enzymes from mesophilic plants, these loop regions moved more freely as temperature increased as indicated by the higher differences in RMSF values for the corresponding aligned residues (Figure 3A). The corresponding loops of CmGLYK moved less than the corresponding aligned residues from AtGLYK as indicated by the near constant RMSF values for this same loop region at different temperatures. These loops presented prime candidates for structure-based recombination due to their greater rigidity in CmGLYK and relative proximity to the Walker A motif. The Walker A motif had low RMSF values in all species simulated at all temperatures, suggesting that fluctuations within this portion of the protein does not explain differential thermostability observed among the GLYK proteins.

**Figure 4.**
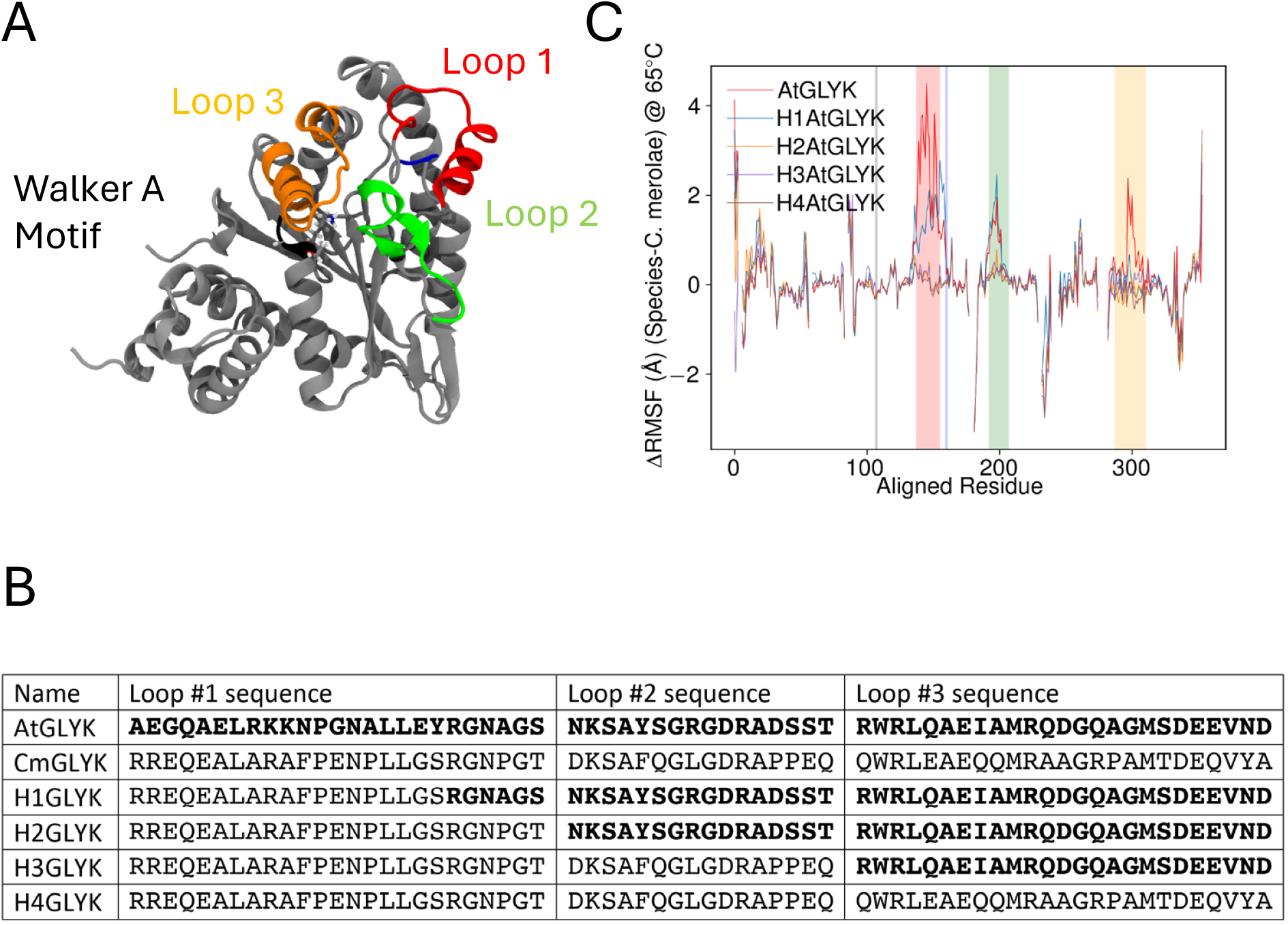
Loops targeted for structure-based recombination and subsequent impact to loop flexibility. A) Loop regions targeted for structure-based recombination indicated by different colors in the enzyme structure with the Walker A Motif (black region) indicated by ball and stick depictions. Loop color corresponds to shaded regions in Figure 3a and 4b. In brief, red is used for the first loop modified in all hybrids, the blue region was chimerized in H2-4AtGLYK, the green region was chimerized in H3-4AtGLYK, and the orange region was only chimerized in H4AtGLYK. B) Amino acid sequences of loop sequences of Arabidopsis and *C. merolae* GLYK proteins and hybrid proteins characterized in this study. Loops are numbered as in A). Amino acid sequences shown in bold match the Arabidopsis GLYK protein while sequences shown in plain text match CmGLYK. C) Change in loop stability at elevated temperature (65°C) measured by comparing RMSF between AtGLYK and its hybrids to the equivalent residues in *C. merolae*.

The three regions from AtGLYK with elevated RMSF values at elevated temperature and relatively close spatial proximity to the ATP binding Walker A motif were selected for replacement with corresponding, more rigid CmGLYK sequences (Figure 4a, 4b, and Supporting Information 3). These various hybrid enzymes were then compared using molecular dynamics to determine which loop replacements resulted in decreased movement on an aligned residue basis. Replacement of a limited portion of Loop 1 (H1AtGLYK) resulted in an enzyme that showed RMSF values closer to AtGLYK than CmGLYK, although there was less motion overall (Figure 4C). When a more extensive portion of Loop 1 was replaced (H2AtGLYK), the Loop 1 region was more rigid and the RMSF was similar to that of CmGLYK. Further replacements of the extended portion of Loop 1 combined with Loop 2 (H3AtGLYK) and Loops 2 and 3 (H4AtGLYK) resulted in no clear changes in the movement of the enzyme from simulation.

Based on these computational results we anticipated that hybrids H2AtGLYK, H3AtGLYK, H4AtGLYK also would have greater thermostability than AtGLYK if these loops were involved in the thermostability of CmGLYK.

### Hybrid enzyme characterization

To evaluate the thermostability of GLYK hybrid enzymes, each hybrid was heterologously expressed in *E. coli*, purified, and characterized. All purified hybrids exhibited glycerate 3-kinase activity. We first characterized the melting temperatures of CmGLYK, AtGLYK, and the four AtGLYK hybrid enzymes using differential scanning fluorimetry with SYPRO Orange (Figure 5A). These analyses revealed that CmGLYK had a melting temperature (T_m_) of 59 °C while AtGLYK had a T_m_ of 53 °C. The hybrids H3AtGLYK and H4AtGLYK with two or three loops replaced correspondingly were more thermostable than the wild-type AtGLYK with their T_m_ increased by 6 °C above that of AtGLYK to a value similar to that of CmGLYK. These findings were consistant when T_m_ was determined using either a fit to a Botzmann equation or by using the negative first derivative (Figure 5a and b).

**Figure 5.**
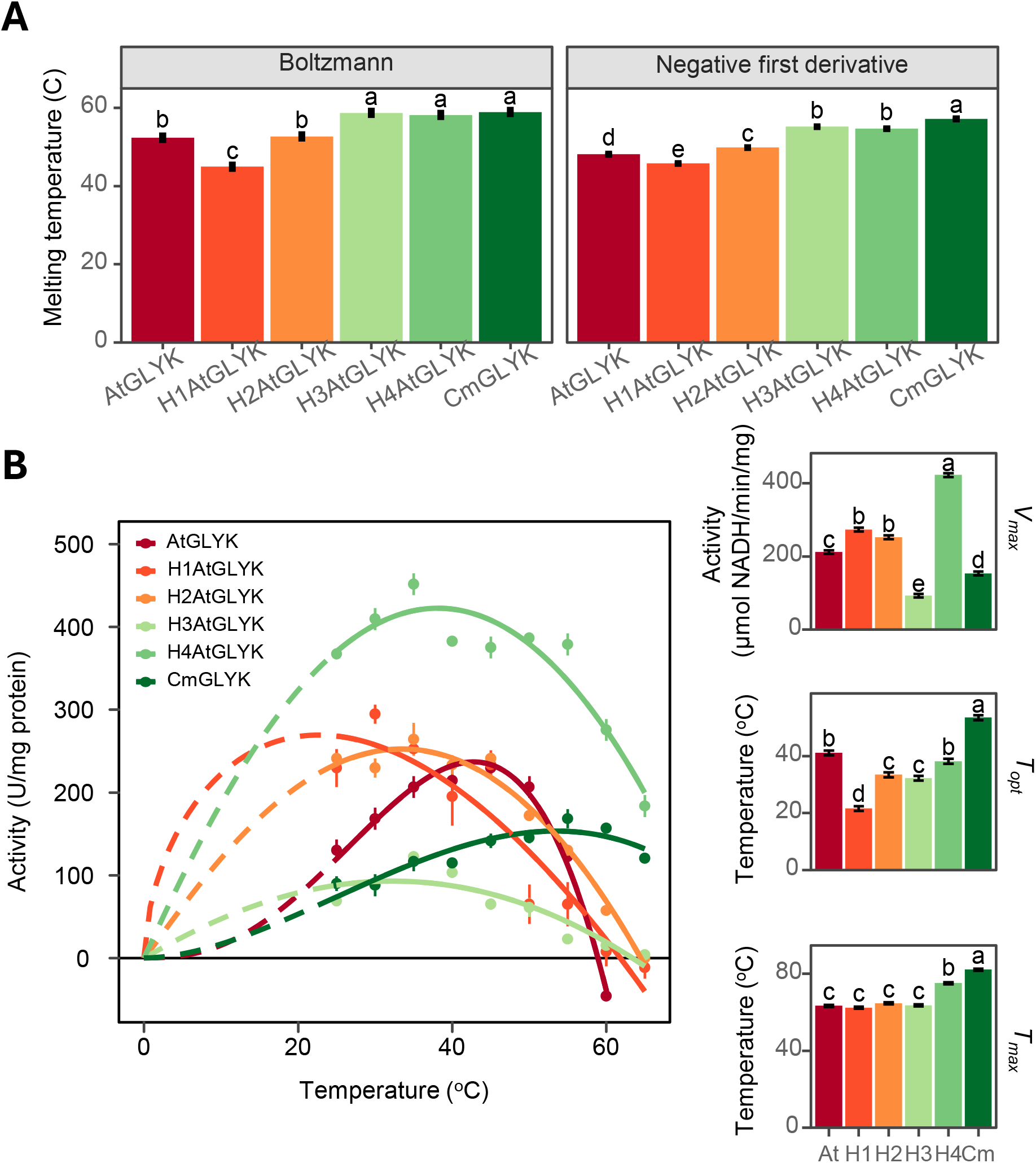
Characterization of thermostability of glycerate kinase from *A. thaliana*, *C. merolae*, and hybrids generated using structure-based recombination. A) Melting temperatures determined from the enzymes using SYPRO orange fluorescence measured as a function of temperature using two independent methods including 1) fitting the sigmoidal portion of the melting curve to the Boltzmann equation, and 2) by determining the maximum melting rate from the negative first derivative of the fluorescence intensity values as a function of temperature (Huynh and Partch, 2015). B) The temperature activity profile for glycerate kinase from *A. thaliana* (AtGLYK), *C. merolae* (CmGLYK) and four hybrid enzymes generated from structure-based recombination (H1AtGLYK, H2AtGLYK, H3AtGLYK, H4AtGLYK) measured using a discontinuous 2-step assay modified from (Kehrer *et al*., 2007) at temperatures from 25°C to 65°C as described in Experimental procedures. Also shown are various parameters determined from the non-linear fit of the temperature response including the maximum activity measured (V_max_), the temperature where V_max_ was measured (T_opt_), and the maximum temperature where activity would be expected (T_max_). Shown are mean ± S.E. n=3. Statistical analysis was performed as described in the Experimental Procedures. Different lowercase letters indicate significant differences between parameters within the same column (p-value < 0.05).

The plot of the first derivative of fluorescence change as a function of temperature resulted in a plot with one minimum for CmGLYK, suggesting a single step denaturation (Supporting Information Figure 4). Multistep denaturation was observed for AtGLYK based on its pattern of the first derivative of fluorescence relative to temperature with multiple, or extended minima. The shape of the first derivative plot for H1AtGLYK and H2AtGLYK resembled that of AtGLYK, whereas the plot for H3AtGLYK and H4AtGLYK resembled the plot of CmGLYK.

To evaluate the temperature response of activity of wild-type and hybrid GLYK enzymes at various temperatures (25-65 °C), the discontinuous GLYK enzyme activity assay was employed. While AtGLYK showed undetectable activity when assayed at 60 °C and 65 °C, the hybrid H4AtGLYK, containing three grafted loops, retained 60% and 40% of its maximal activity, which was comparable to wild-type CmGLYK at these temperatures (Figure 5B).

Interestingly, while the optimal temperature for activity was not increased in the hybrids, the maximum activity as determined from non-linear curve fitting of the entire response was significantly increased in H4AtGLYK.

### Kinetic characteristics of GLYKs

To further characterize GLYK enzymes from different plant species and the recombinant hybrids, the specific activity and kinetic parameters (V_max_, K_m_ and K_cat_) were measured at 30 °C (Table 2). AtGLYK exhibited the highest V_max_ out of all the GLYK homologs tested. The K_m_ values for glycerate for all plant GLYK enzymes were comparable, in the range 0.21 – 0.41 mM, however CmGLYK had a K_m_ approximately 2-4 fold lower of 0.1 mM. A similar trend was observed for the K_m_ for ATP. The K_m_ of plant GLYK proteins for ATP varied from 0.45 to 0.81 mM, whereas the Km of CmGLYK was ∼10-fold lower for this substrate. These data uggest that there may be differences in *in vivo* concentrations of these substrates (glycerate and ATP) in plants as compared to C. *merolae*.

**Table 2.** Kinetic parameters of recombinant glycerate kinase enzymes. Kinetic parameters of glycerate kinase from *Arabidopsis thaliana, Nicotiana tabacum*, *Brassica rapa,* and *Cyanidioschyzon merolae*. Also included are four hybrid enzymes generated using structure- based recombination of *A. thaliana* glycerate kinase using loop regions from the thermophilic *C. merolae* (H1AtGLYK, H2AtGLYK, H3AtGLYK, H4AtGLYK). Kinetics were measured at 30°C and presented as mean ± S.E., n=3. Statistical analysis was performed as described in the Experimental Procedures. Different lowercase letters indicate significant differences between means within the same column according to an ANOVA analysis (p-value < 0.05).

| Enzyme/Hybrid | Glycerate |  |  | ATP |  |  |
| --- | --- | --- | --- | --- | --- | --- |
|  | V<sub>max</sub> (U/mg) | K<sub>m</sub> (mM) | k<sub>cat</sub> (sec<sup>-1</sup>) | V<sub>max</sub> (U/mg) | K<sub>m</sub> (mM) | k<sub>cat</sub> (sec<sup>-1</sup>) |
| *A. thaliana* | 77.99 ± 1.64 b | 0.21 ± 0.017 c | 50.04 ± 1.06 b | 132.09 ± 3.14 a | 0.47 ± 0.025 b | 84.75 ± 2.11 a |
| *N. tabacum* | 37.69 ± 2.16 de | 0.41 ± 0.083 c | 27.59 ± 1.4 cd | 79 ± 1.65 d | 0.81 ± 0.071 a | 57.83 ± 1.11 c |
| *B. rapa* | 31.84 ± 1.65 ef | 0.21 ± 0.043 cd | 22.59 ± 1.06 de | 43.56 ± 1.23 f | 0.45 ± 0.043 bc | 30.9 ± 0.83 f |
| *C. merolae* | 28.86 ± 1.33 f | 0.098 ± 0.02 d | 19.41 ± 0.86 e | 33.15 ± 0.96 g | 0.046 ± 0.0046 d | 22.31 ± 0.64 g |
| H1AtGLYK | 104.73 ± 3.4 a | 0.99 ± 0.087 b | 67.29 ± 2.2 a | 118.76 ± 1.18 b | 0.38 ± 0.012 c | 76.31 ± 0.79 b |
| H2AtGLYK | 86.62 ± 3.92 b | 1.22 ± 0.14 ab | 55.71 ± 2.53 b | 94.49 ± 1.17 c | 0.37 ± 0.015 c | 60.78 ± 0.79 c |
| H3AtGLYK | 52.91 ± 5.24 cd | 4.57 ± 1.11 a | 34.07 ± 3.39 c | 56.56 ± 1.22 e | 0.44 ± 0.032 bc | 36.42 ± 0.82 e |
| H4AtGLYK | 47.89 ± 1.63 c | 1.69 ± 0.2 a | 30.86 ± 1.05 c | 79.62 ± 0.93 d | 0.55 ± 0.023 b | 51.31 ± 0.62 d |

Hybrid enzymes also were assayed to evaluate whether loop replacements impacted kinetic properties. The specific activity for hybrids H2AtGLYK, H3AtGLYK, and H4AtGLYK was approximately 2-fold lower in comparison to AtGLYK but remained more than 2-fold higher than CmGLYK (Table 2). The K_m_ for glycerate for all hybrid enzymes increased relative to AtGLYK, however K_m_ for ATP was not affected. K_cat_ for hybrids H3AtGLYK and H4AtGLYK for glycerate and ATP slightly decreased relative to AtGLYK.

## Discussion

This research highlights the potential for structure-guided protein engineering approaches to improve enzyme thermotolerance in plants. To engineer increased thermostability of GLYK, an essential photorespiratory enzyme, we modeled the three-dimensional structures of GLYK enzymes using the AlphaFold structural prediction tool and used these structures as inputs for classical molecular dynamics simulations. These simulations highlighted candidate loop regions of the AtGLYK enzyme exhibiting high RMSF values that could contribute to instability of this protein at elevated temperatures. We hypothesized that replacing these high RMSF loops with corresponding regions from the more thermotolerant CmGLYK enzyme would increase AtGLYK thermotolerance. To test this hypothesis, we generated a series of hybrid GLYK enzymes consisting of different loop regions from CmGLYK introduced into the AtGLYK backbone. These hybrid enzymes all were functional and, as hypothesized, were more thermostable based on T_m_ measurements (Figure 5). Additionally, H4AtGLYK retained much of its maximal activity at elevated temperatures whereas AtGLYK showed no detectable activity at these temperatures (Figure 5B). Interestingly, H4AtGLYK also had the maximal specific activity out of all the enzymes measured. Based on K_m_ measurements for glycerate and ATP, the grafted loops did not appear to perturb the Walker A motif participating in ATP binding in hybrids and hence did not affect the affinity for ATP, but the grafted loops in hybrids may have indirectly perturbed the binding pocket for glycerate as the K_m_ for glycerate increased 8-10 fold for hybrids H3AtGLYK and H4AtGLYK (Table 2). Another possibility is that the "stiffening" of the enzyme decreased passage of substrate/product in/out of the catalytic site - this may especially be true at 30 °C (temperature used for the assays), and less of an issue at higher temperatures.

Analysis of predicted sequence-based structural surface loops by AlphaFold-assisted molecular dynamics simulations of GLYK proteins from mesophilic plants (*A. thaliana*, *B. rapa*, *N. tabacum*) and a thermophilic alga *C. merolae* uncovered fundamental structural elements that may underlie the increased thermostability of the algal enzyme. The three CmGLYK surface loops selected for grafting were more rigid and compact and fluctuated far less during molecular dynamic simulations than corresponding regions of AtGLYK, suggesting that high fluctuation of these loops in AtGLYK at elevated temperatures may be partially responsible for structural instability and decreased activity at elevated temperatures. These fluctuations were observed as transient unfolding events during simulation (Supporting Information Animations 2 and 3).

While not conclusive, these loop fluctuations may destabilize the protein and enhance unfolding and inactivation of the protein by disrupting the overall structure, or simply by removing the substrate stabilizing interactions needed for efficient catalysis. The correlation of protein stability with lower levels of fluctuations has been reported previously (Tang and Dill, 1998). The knowledge obtained from GLYK as a case study may be more generally applicable for engineering thermotolerance among other enzymes including those not from plants.

Comparing GLYK from plants with the enzyme from *C. merolae* by different approaches such as sequence alignment and amino acid compositional analysis, thermostability, T_m_, kinetics, surface loops structural dynamics, and protein denaturation pattern, demonstrate that these enzymes have different properties that may result from adaptation to different environments. Despite these differences, the AlphaFold predicted structures of these enzymes were nearly identical, with a clearly conserved fold recognized by the neural network architecture underlying AlphaFold. The amino acid composition of CmGLYK provided evidence for adaptation of the enzyme at the level of primary amino acid sequence. A larger proportion of hydrophobic, aromatic or charged amino acids, a decrease in polar amino acids, and substitutions of glycine with alanine and lysine with arginine, have been observed in enzymes originating from thermophilic bacteria and fungi compared to homologs from mesophilic counterparts (de Oliveira *et al*., 2018; Vieille and Zeikus, 2001), however exceptions to these trends have also been reported (Böhm and Jaenicke, 1994). Arginine is a positively charged amino acid, intrinsically carries charge at physiological pH, and is found at higher levels in enzymes of thermophilic and extremophilic bacteria and fungi (Armstrong *et al*., 2016; Li *et al*., 2018; Mrabet *et al*., 1992). Arginine has been identified as an element to enhance protein thermostability, likely by forming salt bridges or hydrogen bonds with neighboring residues (Boucher *et al*., 2022) Additionally, arginine substitutions of lysine improve thermostability of structurally and functionally unrelated enzymes (Li *et al*., 2018; Mrabet *et al*., 1992).

Interestingly, CmGLYK contained a larger proportion of hydrophilic positively charged arginine, but the H3AtGLYK and H4AtGLYK hybrids did not have higher levels of arginine in comparison to AtGLYK and had comparable thermostability to CmGLYK (Table 1, Figure 5).

However, these hybrids contained proline, leucine, glutamic acid, asparagine, and glutamine in quantities close to the levels in CmGLYK, pointing at the potential importance of hydrophobic and hydrophilic polar uncharged amino acid in thermostability for this plant enzyme and highlighting that there is not a single mechanism that provides enzyme thermotolerance.

Elevated temperature increases catalytic rates of enzymes until a temperature threshold is reached, causing thermal denaturation and enzyme inactivation. We followed the kinetics of this denaturation in GLYK through the Sypro Orange thermofluor assays in AtGLYK and CmGLYK. The first derivative plots of the melting curves for these proteins revealed one clearly defined minimum for CmGLYK, consistent with this enzyme possessing a single-phase denaturation process, representing an “all at once” unfolding of this enzyme (Supportive Information Figure 4). By contrast, the first derivative plots for AtGLYK and the hybrids showing at least two minima distributed over a broader temperature stretch indicated a multistep denaturation pattern. Hybrids H1AtGLYK and H2AtGLYK showed multistep denaturation patterns resembling that of AtGLYK, while the denaturation pattern for hybrids H3AtGLYK and H4AtGLYK more closely resembled that of CmGLYK showing a single denaturation minimum. The differential denaturation patterns suggest that loop flexibility evident in AtGLYK molecular dynamics simulations may result in a partially unfolded AtGLYK enzyme possessing less activity.

Mechanistically, H1AtGLYK and H2AtGLYK only alter a single loop, whereas H3AtGLYK and H4AtGLYK feature interloop contacts that further stabilize the structure. Thus, in the case of H1AtGLYK and H2AtGLYK, the identified loop may be the first to unfold at elevated temperature (∼40 °C), decreasing affinity for glycerate and abolishing activity. In contrast, the interloop contact optimizations for H3AtGLYK and H4AtGLYK could mean that unfolding occurs only when the larger catalytic domain breaks down (at around 60 °C). These observations suggested that the different energy required for unfolding grafted loops may contribute to the process of multistep denaturation. We hypothesize that initial steps in denaturation represent the fluctuating loops losing their structure, followed by denaturation of the core domains through a series of transitions.

This study provides an example of applying molecular simulation approaches to engineer functional enzyme hybrids that are more thermostable than mesophilic homologs, however additional studies are needed to understand whether these modifications increase overall plant performance at elevated temperature. For example, the net impact of modifications that increase enzyme thermostability while changing to substrate affinity are not yet clear. It is possible that further efforts to improve thermostability of GLYK could yield a more thermostable hybrid with minimal alteration to its kinetic properties. It also is possible that modifications to enzymes may affect enzyme or pathway regulation *in vivo*. To ensure that enzyme regulatory function stays unchanged, additional enzyme modifications may be needed around key residues in regulatory or active site domains. Further stabilization of the protein core through structure-based approaches and/or by altering amino acid distribution patterns on a larger scale could also be used.

### Experimental procedures

Expression plasmids for plant and *C. merolae* GLYK were generated in the pET28 vector by the U.S. Department of Energy (DOE) Joint Genome Institute (JGI) and provided in NEB T7 express *E. coli* cells. For expression of recombinant GLYK proteins in *E. coli*, sequences encoding chloroplast targeting peptides were removed and an N-terminal His6-tag was included for protein purification.

### Construction of expression plasmids for hybrid AtGLYK

Hybrid GLYK enzymes based on the AtGLYK coding sequence lacking chloroplast targeting peptide and with loop regions replaced with corresponding CmGLYK sequences (Figure 4B and Supporting Information Figure 3) were designed *in silico,* synthesized by GenScript Biotech (Piscataway, NJ, USA), and inserted into the NdeI/XhoI cloning site of the pET28a(+) vector. The resulting plasmids containing hybrid GLYKs with an N-terminal His6 –tag were transformed into NEB T7 express *E. coli* cells using the Mix & Go *E. coli* transformation system (Zymo Research, Irvine, California, USA) according to the manufacturer’s instructions using kanamycin for selection. Glycerol stocks of transformants were stored at -80°C. All plasmid constructs described in this study are available upon request.

### Expression and purification of recombinant GLYKs

Cultures containing 50 ml Luria-Bertani medium supplemented with 50 µg/ml kanamycin were inoculated with 0.5 ml overnight bacterial culture grown in the same medium from a single colony. Cultures were incubated at 37 °C with shaking at 225 rpm until OD_600nm_ reached 0.6-0.8. After cooling cultures to 20 °C, recombinant protein expression was initiated by addition of 0.1 mM IPTG, and the culture was shaken at 20 °C overnight. Cells were harvested by centrifugation at 4,200 *g* for 20 min at 4 °C and frozen in liquid nitrogen. Frozen cells were immediately used for purification or stored at –80 °C. HisPur^TM^ Ni-NTA Spin Columns (Thermo Scientific, Rockford, IL, USA) were used for purification of all recombinant GLYK proteins with some modifications to the protocol (Ni-NTA Spin Kit Handbook, Qiagen). Frozen cells obtained from three 50 ml bacterial cultures were lysed in 4 ml lysis buffer [50 mM Tris-HCl, pH 8.0, 300 mM NaCl, 1 mM EDTA, 9.5 mg/ml lysozyme, 0.1 mg/ml DNAse1, 10 mM CaCl2, 1 tablet proteinase inhibitors cocktail (Roche, Complete Mini, EDTA-free, Mannheim, Germany), 10 mM imidazole, 5% glycerol] on ice for 1 h with gentle shaking, and then sonicated on ice 5x10 sec using a sonicator (Branson Sonifire 150, Danbury, CT, USA) at high frequency, with 10 sec intervals for cooling. The soluble proteins were collected as the supernatant after centrifugation at 19,800 g for 20 min at 4 °C. The purification of the recombinant protein was performed basically as described by the manufacturer of the HisPur Ni-NTA spin columns (ThermoScientific, Rockford, IL, USA), with the following modifications: 50 mM Tris-HCl, pH 8.0, 300 mM NaCl, containing 10 mM, 20 mM, 500 mM imidazole was used for binding, washing and elution correspondingly. Binding of recombinant protein to the affinity matrix was performed by addition of 600 μl of soluble protein to a 0.2 ml Ni-NTA column followed by centrifugation for 2 min at 50 g at 4°C, this process was repeated until all soluble protein had been loaded followed by three washes using 50 mM Tris-HCl, pH 8.0, 300 mM NaCl, 20 mM imidazole. Recombinant His-tagged GLYK was eluted twice using 300 μl of elution buffer containing 500 mM imidazole, and combined eluates were dialyzed twice against 250 ml (50 mM Tris-HCl, pH 8.0, 300 mM M NaCl) at 4 °C, one hour each, then overnight against 1000 ml 50 mM Tris-HCl, pH 8.0, 300 mM M NaCl, 5% glycerol.

Protein concentrations were estimated using NanoDrop and/or Quick Start Bradford reagent (BioRad, Hercules, CA, USA) with bovine serum albumin as a standard. Sterile glycerol was added to purified protein to final 10%, and the protein solution was stored in aliquots at -80 °C.

### SDS-PAGE Analysis

*E. coli* crude soluble fractions, purified recombinant proteins, and protein fractions obtained during purification procedures were separated by SDS-PAGE using pre-cast 10% [w/v] acrylamide gels (BioRad, Hercules, CA, USA) and detected by Coomassie blue staining (Laemmli, 1970).

### Glycerate 3-kinase activity assay

Glycerate kinase activity was measured by coupling the formation of 3-phosphoglycerate to NADH oxidation with the use of coupling enzymes in a 96- well microplate format (Gregory *et al*., 2023). NADH consumption by the coupling enzymes was monitored by absorbance measured continuously with a plate reader at 340 nm during the first two minutes of kinase reaction. Briefly, reaction mixtures with a total volume of 200 μl, contained 192 µl reaction buffer (50 mM HEPES, pH 7.8, 10 mM MgCl_2_, 60 mM KCl, 5 mM ATP, 5 mM creatine phosphate, 0.2 mM NADH, 4 μl (50 ng) purified GLYK, and 4 µl coupling enzymes (22.5 U ml^-1^ 3-phosphoglycerate kinase, 12.5 U ml^-1^ creatine phosphokinase, 20 U ml^-1^ glyceraldehyde-3-phosphate dehydrogenase, 20 U mL-1 glycerol-3-phosphate dehydrogenase, 56 U ml^-1^ triose-phosphate isomerase, all from Sigma-Aldrich, St. Louis, MO, USA). The reaction was initiated by addition 2 μl of 500 mM glycerate using a 96-well plate replicator.

Kinetic parameters, K_m_ and V_max_, were determined by fitting the response of enzyme activity to different concentrations of substrate using standard Michaelis-Menton kinetics for a one-substrate enzyme-catalyzed reaction. Concentrations of substrate ranged from 0–10 mM for glycerate and or 0-5 mM for ATP. Code for this fitting procedure is available at https://github.com/PerennialDr/MichaelisMenten_fit and implemented as a user-friendly fitting app at https://perennialdr.shinyapps.io/MichaelisMentenFit/, . The V_max_ value was used to calculate the turnover number k_cat_. All measurements were performed in triplicate. The activity was expressed as μmoles NADH utilized per minute per milligram of protein.

### GLYK activity thermostability assays

Two approaches were employed to characterize GLYK activity thermostability. For the first approach, aliquots of GLYK in 50 mM HEPES, pH 7.5, containing 25 μg/ml GLYK, were treated at 4 °C, 25 °C, 30 °C, 35 °C, 40 °C, and 45 °C for 0, 15, 30, and 60 minutes. GLYK activity was measured at 30 °C immediately after each treatment. For the second approach to determine the optimal reaction temperature, a discontinuous 2-step assay modified from (Kehrer *et al*., 2007) was used to measure the activity of GLYK at temperatures from 25°C to 65°C. In the first step of this assay, 388 µl reaction buffer containing 50 mM HEPES, pH 7.5, 10 mM MgCl_2_, 60 mM KCl, 5 mM ATP, 5 mM D-glyceric acid, was warmed to the desired temperature in a heating block. The reaction was initiated by adding 8 µl 50 mM HEPES, pH 7.5, containing 200 ng recombinant GLYK. After 1 min the reaction was terminated by transferring the tube into a heating block at 95 °C for 5 min. The second step was performed in a 96-well microplate at 30 °C. 96 µl reaction buffer 2 containing 50 mM HEPES, pH 7.5, 10 mM MgCl_2_, 60 mM KCl, 15 mM NADH, 2 mM ATP, 10 mM creatine phosphate, was dispensed per well into a plate pre-warmed to 30 °C. 100 µl of the reaction mix obtained at the first step was added and the initial optical density at 340 nm was observed for 2 min using the microplate reader. The reaction was initiated by adding 4 µl of coupling enzymes (22.5 U ml^-1^ 3- phosphoglycerate kinase, 12.5 U ml^-1^ creatine phosphokinase, 20 U ml^-1^ glyceraldehyde-3- phosphate dehydrogenase, 20 U mL^-1^ glycerol-3-phosphate dehydrogenase, 56 U ml^-1^ triose- phosphate isomerase), and the optical density at 340 nm was monitored for 2 min using the microplate reader. Activity was calculated using the difference between the initial values of optical density and optical density obtained after addition of coupling enzymes, subtracting the background optical density values contributed by coupling enzymes only.

### Melting temperature assay

For determining T_m_, differential scanning fluorimetry was employed using SYPRO Orange (Huynh and Partch, 2015). For these assays, fluorescence measurements were collected from 30 μl reactions in white, 0.2 ml PCR tubes with optical flat caps (Bio-Rad Laboratories, Hercules, California, USA) containing 12.5 mM Na-phosphate (pH 7.5), 15x SYPRO Orange (Sigma, Sigma-Aldrich, St. Louis, MO, USA), and 15 μg of His6- tagged recombinant enzyme (Ginnard *et al*., 2022). Starting at 20 °C, samples were warmed to 95 °C at 1 °C per min with readings collected every 15 seconds using a 7500 Fast Real-Time PCR System equipped with a ROX filter (Applied Biosystems, Thermo Fisher Scientific, Waltham, MA USA). Eight replicates were conducted for each sample with appropriate controls for water and buffer. Raw fluorescence measurements exported to the Design and Analysis software (Applied Biosystems, Thermo Fisher Scientific, Waltham, MA USA) were used to determine T_m_ by two independent methods with the use of R. The first method fitted the sigmoidal portion of the melting curve to the Boltzmann equation, and the second one determined the maximum melting rate from the negative first derivative of the fluorescence intensity values as a function of temperature (Huynh and Partch, 2015). T_m_ estimated using both techniques generally agreed closely, and T_m_ values estimated by both techniques are presented.

### AlphaFold structure prediction

Protein sequences for GLYK were obtained from a variety of sources. For *A. thaliana,* AtGLYK was translated from the coding regions provided from The Arabidopsis Information Resource database (ATG80380). The *C. merolae* GLYK sequence was deduced from the *C. merolae* genome project v3 as entry CMK141C (Matsuzaki *et al*., 2004; Nozaki *et al*., 2007). *B. rapa* GLYK-1 was determined based on homology to AtGLYK through a blast search in Phytozome (Goodstein *et al*., 2012) against the *Brassica rapa* FPsc v1.3 genome and was the coding region for transcript Brara.G03734. *N. tabacum* was determined by a translated nucleotide blast resulting in NCBI reference sequence NCBI XP_016437660. Sequences of mature GLYKs used for structural predictions were generated by removing the predicted chloroplast targeting signal as determined from homology to the chloroplast transit peptide in *A. thaliana* (Supporting Information Figure 5). These sequences were used as inputs for AlphaFold 2.1.1 (Jumper *et al*., 2021), using the published defaults and a maximum template date of November 1^st^, 2021. To generate sequences of GLYK hybrid proteins, a sequence alignment was performed using ClustalX (Larkin *et al*., 2007), and protein sequences near high RMSF regions of AtGLYK were replaced with corresponding *C. merolae* sequences (Supporting Information Figure 3). These novel hybrid sequences were used as inputs for AlphaFold 2.1.1 based on the same methodology but using a maximum template date of October 1^st^, 2022. To facilitate structure comparison between different sequences, we aligned the proteins using US-align (Zhang *et al*., 2022).

### Molecular dynamics simulations and analyses

AlphaFold structural predictions of GLYK variants were solvated in a water cube with a 120 Å edge length using the solvate plugin of Visual Molecular Dynamics (VMD) 1.9.4a57 (Humphrey *et al*., 1996). Once solvated, selected water molecules were replaced with ions to achieve a 150 mM NaCl concentration using the autoionize plugin of VMD. Preparation simulations were carried out in two stages, a brief minimization using NAMD 2.14, followed by 10 ns of equilibration simulation using NAMD 3.0a9 to maximize performance using the GPU-resident integrator (Phillips *et al*., 2020). The equilibration simulation was carried out in the constant pressure and temperature ensemble, using an isotropic Langevin barostat (Feller *et al*., 1995) set at 1 atmosphere and a Langevin thermostat (Feller *et al*., 1995; Kubo, 1966) set to 298K (25°C). As is standard for the CHARMM36m force field for proteins (Huang *et al*., 2017) together with the TIP3 water model (Jorgensen *et al*., 1983), non-bonded terms were switched at 10 Å with a 12 Å cutoff. Long range electrostatic interactions were treated with the particle mesh Ewald method (Darden *et al*., 1993) using a 1.2 Å grid spacing. 2 fs time steps were enabled by using the SETTLE algorithm (Miyamoto and Kollman, 1992) during equilibration. After the 10 ns of equilibrium simulation, each simulation was transferred into a GROMACS compatible format using TopoGromacs (Vermaas *et al*., 2016). Using GROMACS 2022 (Abraham *et al*., 2015), the equilibrated configuration was extended by 200 ns in 5 statistical replicates in a constant volume and temperature ensemble, for a total of 1 μs of sampling for every combination of temperature and protein identity. Temperature was controlled with a velocity rescaling thermostat (Bussi *et al*., 2007), set to 298K (25° C), 318K (45° C), or 338K (65°C) to collect statistics at multiple temperatures. As with NAMD, particle mesh Ewald (Essmann *et al*., 1995) was used for long-range electrostatics with a 1.2 Å grid spacing. LINCS (Hess *et al*., 1997) was used to fix hydrogen bond lengths and enable 2 fs time steps.

Analysis was performed via scripts that leverage the python bindings built into VMD (Humphrey *et al*., 1996), facilitating the use of the numpy (Harris *et al*., 2020). The quantities of interest were the RMSD, which measures the deviation to a selected reference structure, and the RMSF, which measures the fluctuation away from the average structure. The RMSD reference in this case was chosen to be the first frame of the 200 ns production simulations and included all sidechain motion within the RMSD. RMSF was measured after aligning the structure to minimize the RMSD on a per-residue basis using the RMSF per residue function built into the VMD atom selection interface, which averages together the atomic RMSF over a full residue to return a single value. Analysis, inputs, and selected simulation outputs are available from Zenodo (10.5281/zenodo.10794947).

### Statistical analysis

Non-linear models were used to test differences in enzyme kinetics and enzyme thermal properties. To test for differences in V_max_, K_m_ and K_cat_ among enzymes for each substrate separately, enzyme was included as a fixed effect in the model within the gnls function (Pinheiro JC, Bates DM, 2000) . The same statistical analysis was performed to test the differences in GLYK activity in thermostability experiments. Enzyme parameters were compared using the emmeans function within the emmean R package (Lenth, R. *et al*., 2018). Differences between values were considered significant at p-value < 0.05.

## Supporting information

Animation 1

Animation 2

Animation 3

## Authorship

BJW, AHL, JVV, and LVR conceived the research idea for this study with the experimental work performed by LVR. MTN provided data analysis of the experimental work. JVV led the molecular dynamics simulation work with assistance from DS and AA. LVR, DS, AHL, MTN, JVV, and BJW contributed to the writing of the manuscript.

## Acknowledgements

We thank Dr. Dave Bauer from ThermoFisher Scientific for fruitful discussions about conducting melting temperature assays. NEB T7 express *E. coli* clones containing expression plasmids for plant and *C. merolae* GLYK proteins were provided by the U.S. Department of Energy (DOE) Joint Genome Institute, a DOE Office of Science User Facility (Contract No. DE-AC02-05CH11231). This work was funded by the Division of Chemical Sciences, Geosciences, and Biosciences, Office of Basic Energy Sciences of the United States Department of Energy (DE-FG02-91ER20021) and awards from the National Science Foundation Division of Integrative Organismal Systems (2030337 and 2030295). This work was supported in part through computational resources and services provided by the

Institute for Cyber-Enabled Research at Michigan State University. This work used Delta at the National Center for Supercomputing Applications (NCSA) through allocation BIO210061 from the Advanced Cyberinfrastructure Coordination Ecosystem: Services & Support (ACCESS) program, which is supported by National Science Foundation grants #2138259, #2138286, #2138307, #2137603, and #2138296. This material is based upon work supported in part by the Great Lakes Bioenergy Research Center, U.S. Department of Energy, Office of Science, Biological and Environmental Research Program under Award Number DE-SC0018409. The authors declare no conflicts of interest.

## Supporting Information

**Supporting Information Figure 1. Multiple sequence alignment of amino acid sequences of mature GLYKs from different species.** The conserved amino acids are marked with blue color. The canonical Walker A motif, APQGCGKT for plant GLYK and CPQGGGKT for CmGLYK, is enclosed in a red dashed box. Alignment was produced using Jalview. Also shown are the calculated identity and similarity % to AtGLYK

**Supporting Information Figure 2.** Atomically detailed structural models of GLYKs generated with AlphaFold shown as a structural alignment with different colors representing various species as indicated in the figure legend.

**Supporting Information Figure 3.** AtGLYK hybrids amino acid sequence**s.** Red indicates equivalent sequences from CmGLYK and various colors represent loop regions modified using structure-based recombination for each hybrid enzymes (H1GLYK, H2GLYK, H3GLYK, and H4GLYK)

**Supporting Information Figure 4.** Plot of the first derivative of fluorescence relative to temperature for AtGLYK, CmGLYK and four AtGLYK hybrids (H1AtGLYK, H2AtGLYK, H3AtGLYK, H4AtGLYK). Melting curves were generated for eight replicates for each sample as described in the Experimental procedures, and the first derivative plots were generated using Design and Analysis software 2.6.0.

**Supporting Information Figure 5.** GLYK sequences without targeting peptide. In red – Walker A motif.

**Supporting Information Animation 1.** Animation of atomically detailed structural models of GLYKs generated with AlphaFold shown as a structural alignment with different colors representing various species as indicated in the figure legend for Supporting Information Figure 2.

**Supporting Information Animation 2.** Animation of AtGLYK simulated at 25°C, retaining the coloration of figure 4A. The loops make only small motions at this temperature.

**Supporting Information Animation 3.** Animation of AtGLYK simulated at 65°C, retaining the coloration of figure 4A. The loops make large movements at this temperature.

