## Supplemental Figures for "Advancing thermostability of the key photorespiratory enzyme glycerate 3-kinase by structure-based recombination"

### Slide 1
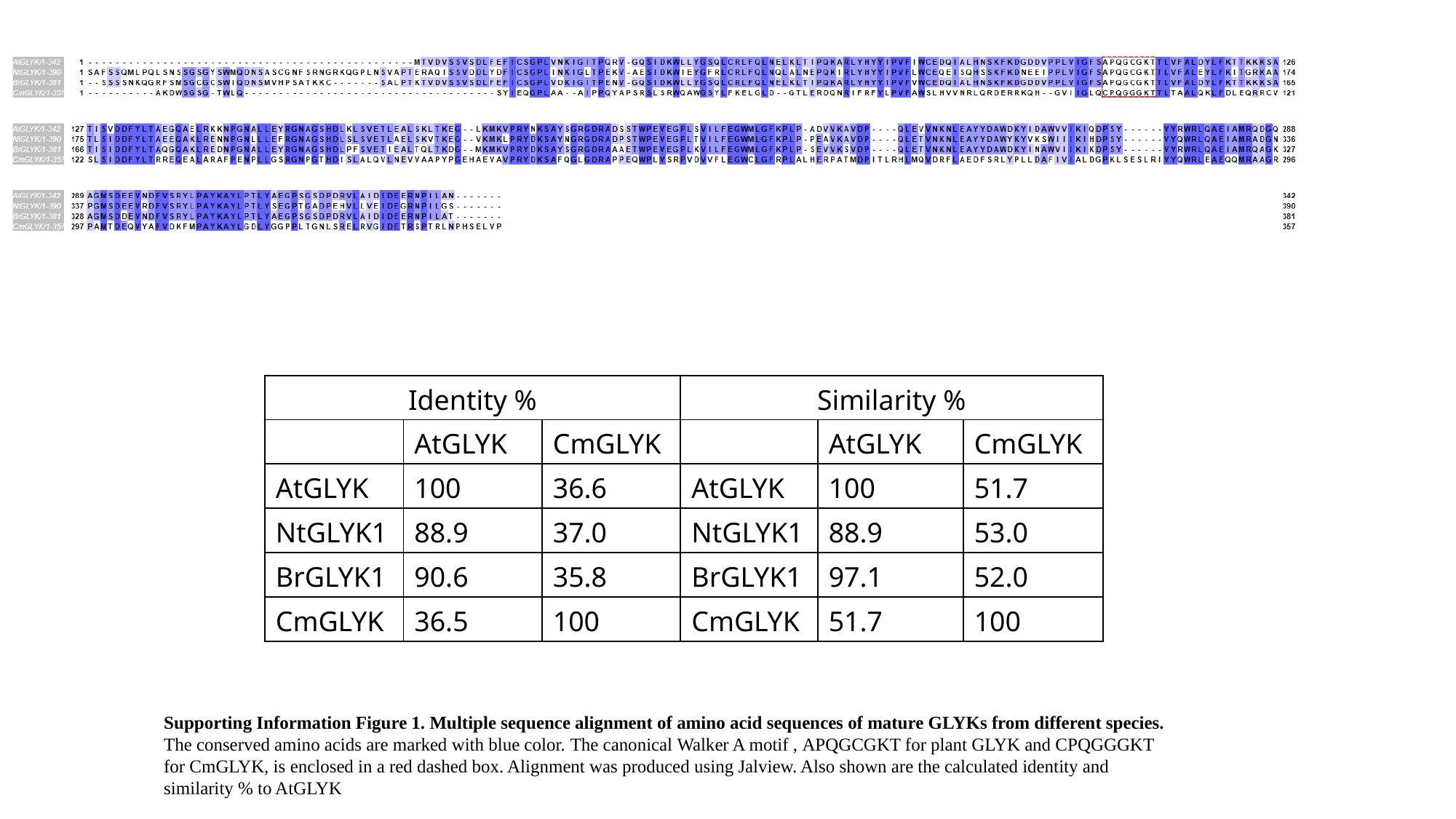

| Identity % | Identity % | | Similarity % | Similarity % | |
| --- | --- | --- | --- | --- | --- |
| | AtGLYK | CmGLYK | | AtGLYK | CmGLYK |
| AtGLYK | 100 | 36.6 | AtGLYK | 100 | 51.7 |
| NtGLYK1 | 88.9 | 37.0 | NtGLYK1 | 88.9 | 53.0 |
| BrGLYK1 | 90.6 | 35.8 | BrGLYK1 | 97.1 | 52.0 |
| CmGLYK | 36.5 | 100 | CmGLYK | 51.7 | 100 |
Supporting Information Figure 1. Multiple sequence alignment of amino acid sequences of mature GLYKs from different species.
The conserved amino acids are marked with blue color. The canonical Walker A motif , APQGCGKT for plant GLYK and CPQGGGKT for CmGLYK, is enclosed in a red dashed box. Alignment was produced using Jalview. Also shown are the calculated identity and similarity % to AtGLYK

### Slide 2
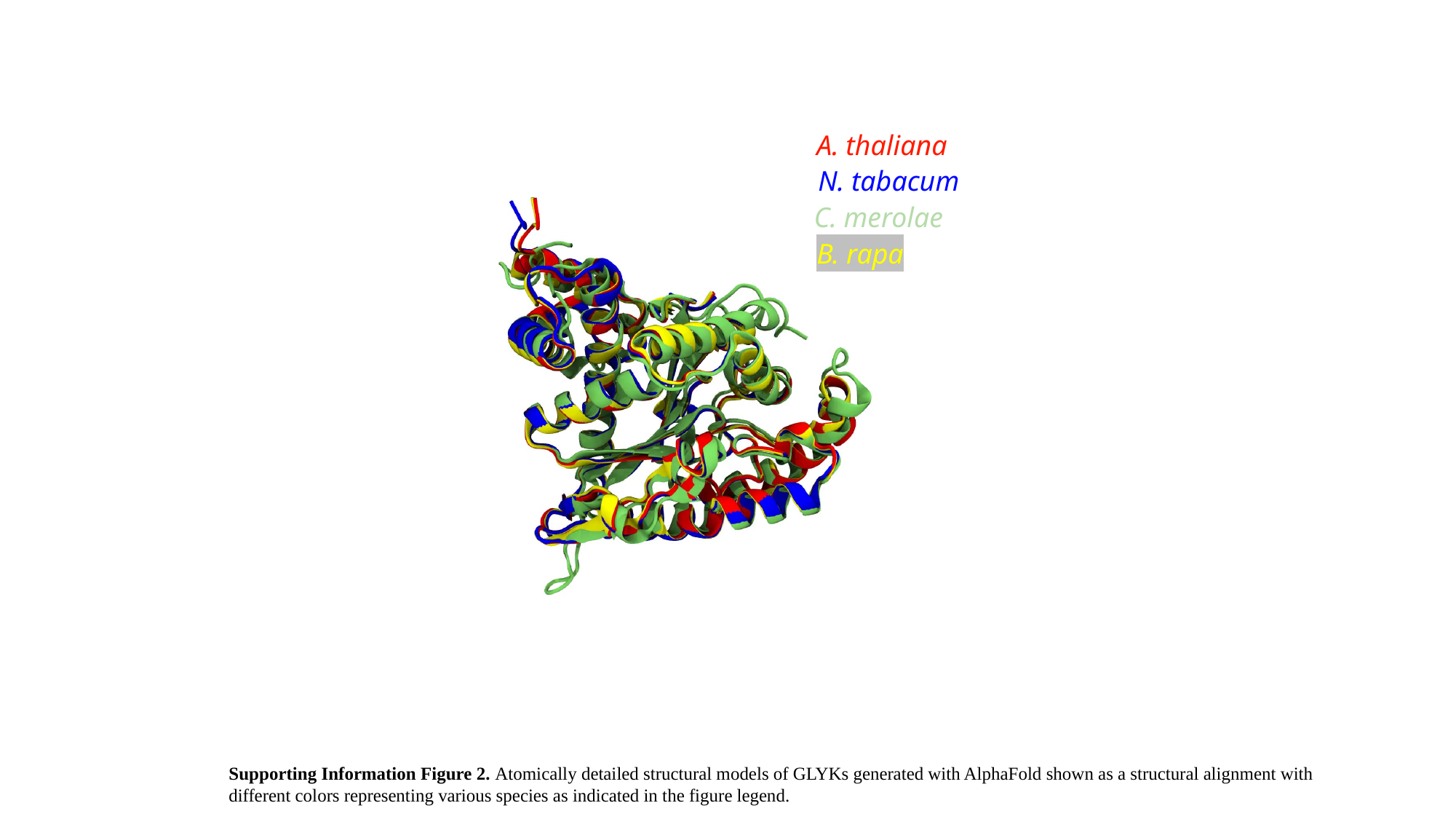

A. thaliana
N. tabacum
C. merolae
B. rapa
Supporting Information Figure 2. Atomically detailed structural models of GLYKs generated with AlphaFold shown as a structural alignment with different colors representing various species as indicated in the figure legend.

### Slide 3
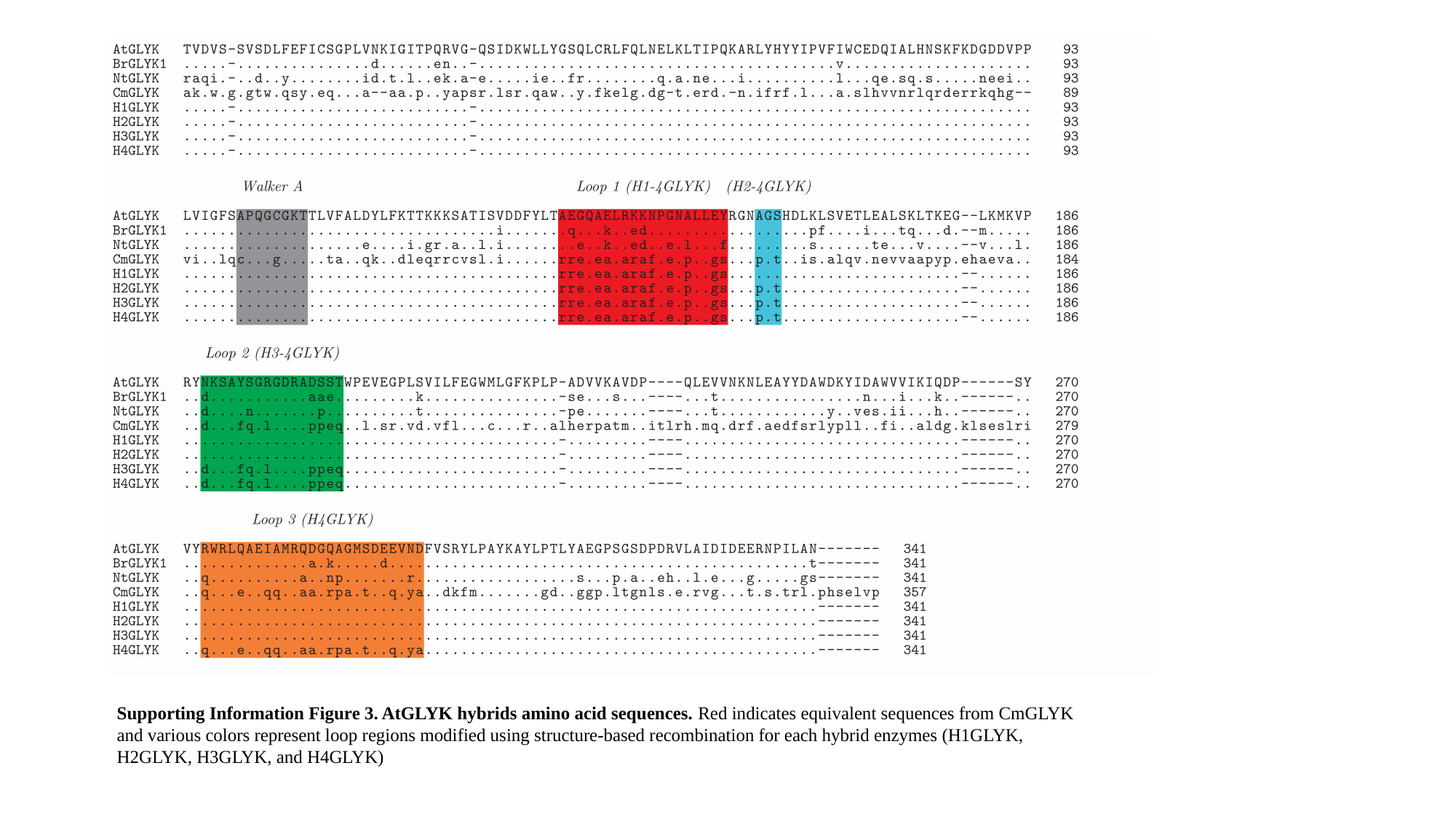

Supporting Information Figure 3. AtGLYK hybrids amino acid sequences. Red indicates equivalent sequences from CmGLYK and various colors represent loop regions modified using structure-based recombination for each hybrid enzymes (H1GLYK, H2GLYK, H3GLYK, and H4GLYK)

### Slide 4
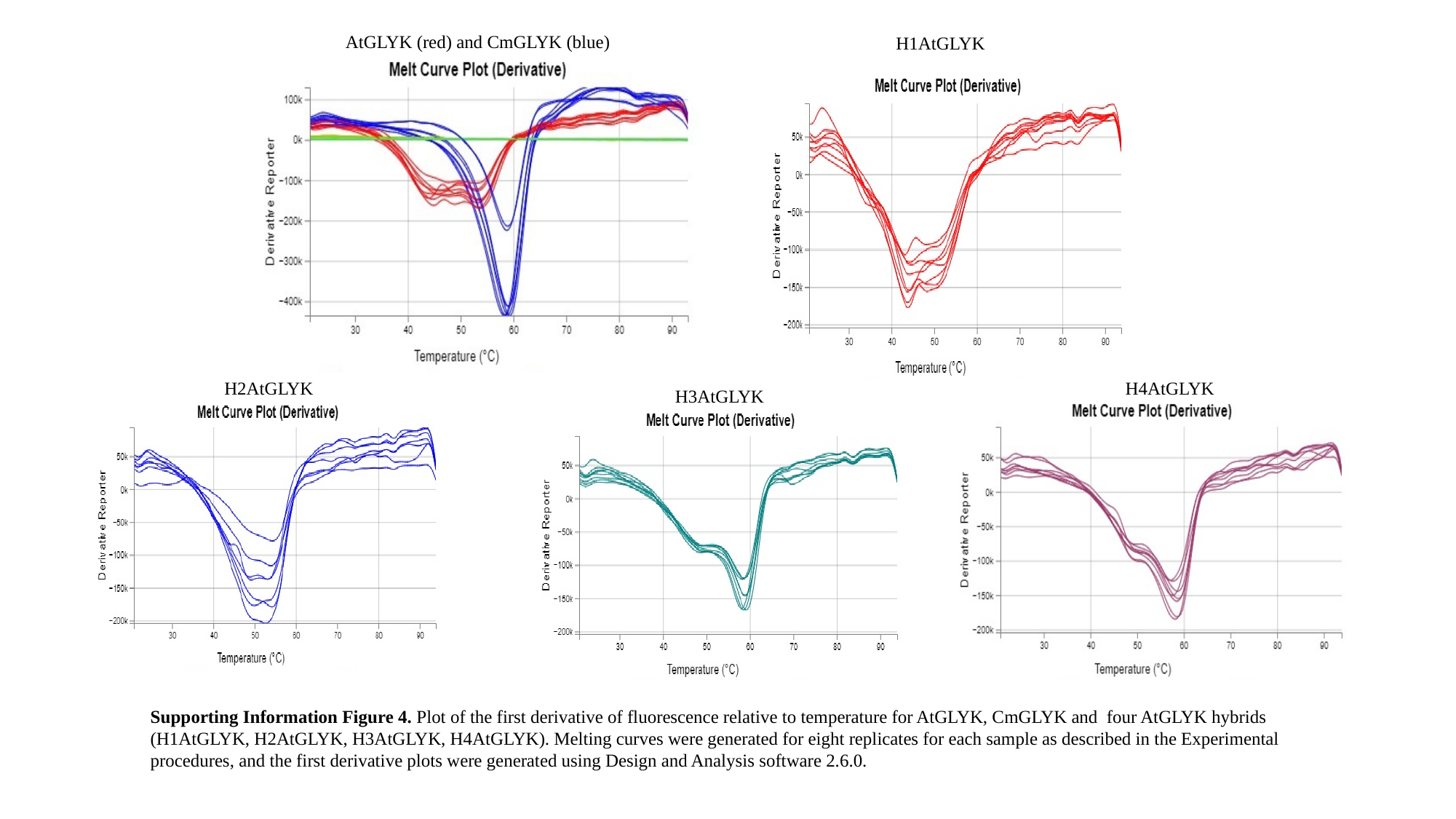

H1AtGLYK
AtGLYK (red) and CmGLYK (blue)
H2AtGLYK
H4AtGLYK
H3AtGLYK
Supporting Information Figure 4. Plot of the first derivative of fluorescence relative to temperature for AtGLYK, CmGLYK and four AtGLYK hybrids (H1AtGLYK, H2AtGLYK, H3AtGLYK, H4AtGLYK). Melting curves were generated for eight replicates for each sample as described in the Experimental procedures, and the first derivative plots were generated using Design and Analysis software 2.6.0.

### Slide 5
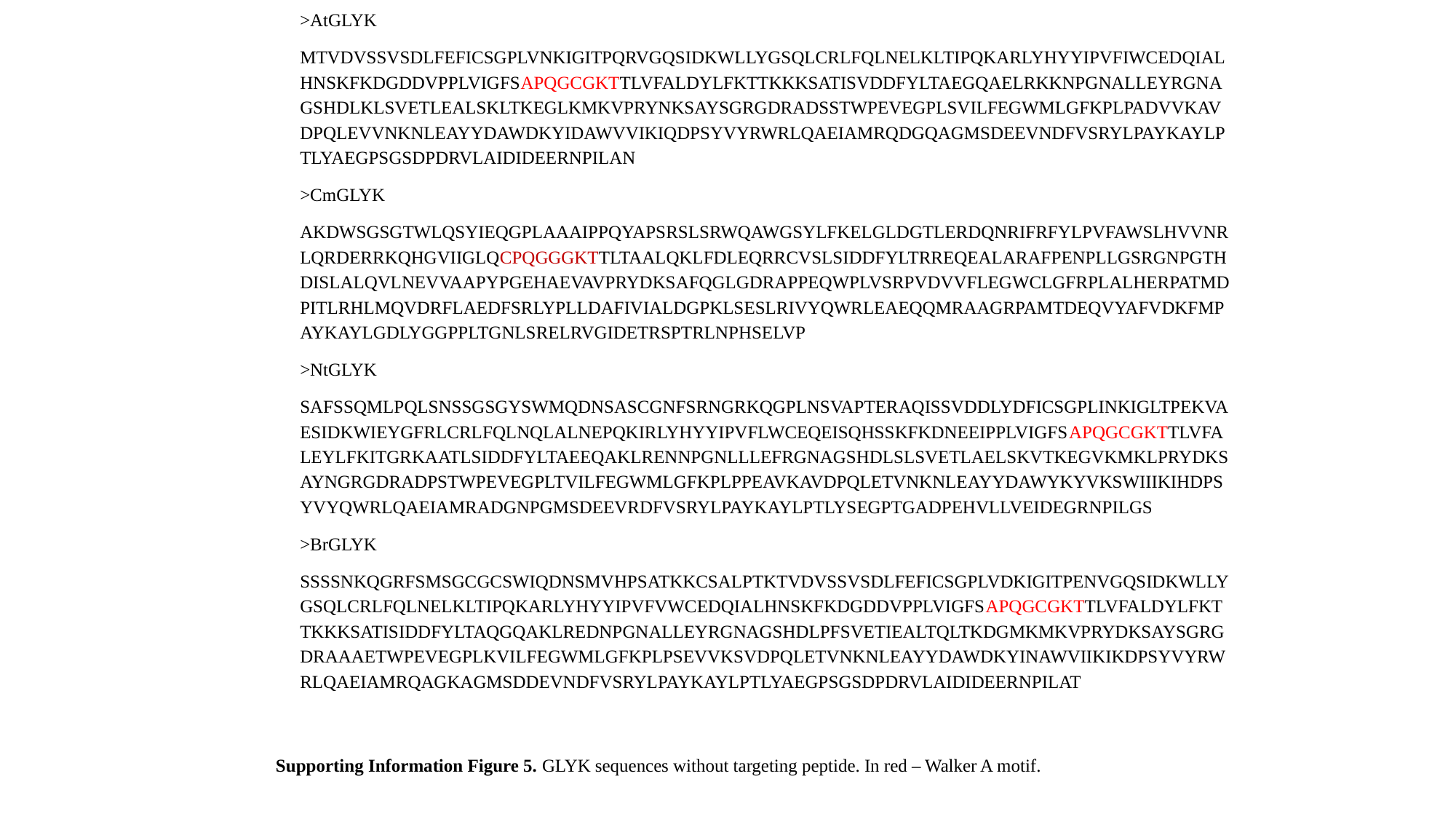

>AtGLYK
MTVDVSSVSDLFEFICSGPLVNKIGITPQRVGQSIDKWLLYGSQLCRLFQLNELKLTIPQKARLYHYYIPVFIWCEDQIALHNSKFKDGDDVPPLVIGFSAPQGCGKTTLVFALDYLFKTTKKKSATISVDDFYLTAEGQAELRKKNPGNALLEYRGNAGSHDLKLSVETLEALSKLTKEGLKMKVPRYNKSAYSGRGDRADSSTWPEVEGPLSVILFEGWMLGFKPLPADVVKAVDPQLEVVNKNLEAYYDAWDKYIDAWVVIKIQDPSYVYRWRLQAEIAMRQDGQAGMSDEEVNDFVSRYLPAYKAYLPTLYAEGPSGSDPDRVLAIDIDEERNPILAN
>CmGLYK
AKDWSGSGTWLQSYIEQGPLAAAIPPQYAPSRSLSRWQAWGSYLFKELGLDGTLERDQNRIFRFYLPVFAWSLHVVNRLQRDERRKQHGVIIGLQCPQGGGKTTLTAALQKLFDLEQRRCVSLSIDDFYLTRREQEALARAFPENPLLGSRGNPGTHDISLALQVLNEVVAAPYPGEHAEVAVPRYDKSAFQGLGDRAPPEQWPLVSRPVDVVFLEGWCLGFRPLALHERPATMDPITLRHLMQVDRFLAEDFSRLYPLLDAFIVIALDGPKLSESLRIVYQWRLEAEQQMRAAGRPAMTDEQVYAFVDKFMPAYKAYLGDLYGGPPLTGNLSRELRVGIDETRSPTRLNPHSELVP
>NtGLYK
SAFSSQMLPQLSNSSGSGYSWMQDNSASCGNFSRNGRKQGPLNSVAPTERAQISSVDDLYDFICSGPLINKIGLTPEKVAESIDKWIEYGFRLCRLFQLNQLALNEPQKIRLYHYYIPVFLWCEQEISQHSSKFKDNEEIPPLVIGFSAPQGCGKTTLVFALEYLFKITGRKAATLSIDDFYLTAEEQAKLRENNPGNLLLEFRGNAGSHDLSLSVETLAELSKVTKEGVKMKLPRYDKSAYNGRGDRADPSTWPEVEGPLTVILFEGWMLGFKPLPPEAVKAVDPQLETVNKNLEAYYDAWYKYVKSWIIIKIHDPSYVYQWRLQAEIAMRADGNPGMSDEEVRDFVSRYLPAYKAYLPTLYSEGPTGADPEHVLLVEIDEGRNPILGS
>BrGLYK
SSSSNKQGRFSMSGCGCSWIQDNSMVHPSATKKCSALPTKTVDVSSVSDLFEFICSGPLVDKIGITPENVGQSIDKWLLYGSQLCRLFQLNELKLTIPQKARLYHYYIPVFVWCEDQIALHNSKFKDGDDVPPLVIGFSAPQGCGKTTLVFALDYLFKTTKKKSATISIDDFYLTAQGQAKLREDNPGNALLEYRGNAGSHDLPFSVETIEALTQLTKDGMKMKVPRYDKSAYSGRGDRAAAETWPEVEGPLKVILFEGWMLGFKPLPSEVVKSVDPQLETVNKNLEAYYDAWDKYINAWVIIKIKDPSYVYRWRLQAEIAMRQAGKAGMSDDEVNDFVSRYLPAYKAYLPTLYAEGPSGSDPDRVLAIDIDEERNPILAT
Supporting Information Figure 5. GLYK sequences without targeting peptide. In red – Walker A motif.
